# Quantifying the degree of sharing of genetic and non-genetic causes of gene expression variability across four tissues

**DOI:** 10.1101/053355

**Authors:** Alfonso Buil, Ana Viñuela, Andrew A. Brown, Matthew N. Davies, Ismael Padioleau, Deborah Bielser, Luciana Romano, Daniel Glass, Paola Di Meglio, Kerrin S. Small, Timothy D. Spector, Emmanouil T. Dermitzakis

## Abstract

Gene expression can provide biological mechanisms which underlie genetic associations with complex traits and diseases, but often the most relevant tissue for the trait is inaccessible and a proxy is the only alternative. Here, we investigate shared and tissue specific patterns of variability in expression in multiple tissues, to quantify the degree of sharing of causes (genetic or non-genetic) of variability in gene expression among tissues. Using gene expression in ~800 female twins from the TwinsUK cohort in skin, fat, whole blood and lymphoblastoid cell lines (LCLs), we identified 9166 significant cis-eQTLs in fat, 9551 in LCLs, 8731 in skin and 5313 in blood (1% FDR). We observed up to 80% of cis-eQTLs are shared in pairs of tissues. In addition, the cis genetic correlation between tissues is > 90% for 35% of the genes, indicating for these genes a largely tissue-shared component of cis regulation. However, variance components show that cis genetic signals explain only a small fraction of the variation in expression, with from 67–87% of the variance explained by environmental factors, and 53% of the genetic effects occurring in trans. We observe a trans genetic correlation of 0 for all genes except a few which show correlation between fat and skin expression. The environmental effects are also observed to be entirely tissue specific, despite related tissues largely sharing exposures. These results demonstrate that patterns of gene expression are largely tissue specific, strongly supporting the need to study higher order regulatory interactions in the appropriate tissue context with large samples sizes and diversity of environmental contexts.

---

Gene expression is an intermediate phenotype between the genome and disease manifestation and it can be used to give a biological and mechanistic interpretation of GWAS signals. However, regulatory control of gene expression is often tissue-specific [1] . It remains an open question how much about the regulatory landscape of one tissue can be inferred given a comprehensive characterization of a different tissue [2]. As disease and phenotypic variation is often linked to inaccessible tissues in living individuals, answering this question could have great impact on our ability to use expression QTLs from heterologous tissues to interpret GWAS signals. Previous studies have compared the effect of common cis-eQTLs across different tissues [1] [3] [4] [5]. However, these common eQTLs represent a small fraction of the genetic effects that regulate gene expression [4]. In this study we provide global measures of the relative contribution of shared and tissue-specific genetic effects on gene expression. We show that genetic regulation of gene expression is largely driven by tissue-specific trans effects. We also show that, though genetic variants in cis tend to be active in more than one tissue, in many cases these genetic variants are affected by tissue specific modulators, which increase the divergence between tissues. These results have implications on the use of eQTLs found in commonly collected tissue types as a tool to interpret the biological function of GWAS signals and our understanding of the biological processes underlying disease development.

Here we analyzed a sample of around 800 female monozygotic (MZ) and dizygotic (DZ) twins from the TwinsUK cohort. For each sample we sequenced the mRNA fraction of the transcriptome in four tissues (fat, skin, lymphoblastoid cell lines (LCLs) and whole blood) and analyzed them with genotypes imputed into the 1000 Genomes Phase 1 panel [6]. To identify regulatory genetic variants affecting gene expression we performed three analyses: expression quantitative trait loci (eQTLs), alternative splicing QTLs (asQTL) and allelic specific expression (ASE). Using a linear regression approach with SNPs in a 1Mb window each side of the TSS for each gene we identified 9166 significant cis-eQTLs in fat, 9551 in LCLs, 8731 in skin and 5313 in blood (1% FDR) (Supplementary Figs. 1 and 2). Genetic variation may also affect gene expression by modifying mRNA splicing processes. Therefore, we calculated the association between cis SNPs and the individual frequencies of exon-exon links and identified between 1566 and 4104 alternative splicing QTLs (asQTLs) per tissue (Supplementary Fig. 3) using ALTRANS [7] . The identification of allele specific expression (ASE) sites has been previously described in Buil et al (2015) and reported that 9.5%, 9.3% and 9.1% of the exonic heterozygous sites had a significant (10% FDR) ASE effect in fat, LCLS and skin. In total, we found thousands of cis regulatory genetic variants affecting gene expression in the four tissues.

To investigate the level of shared cis-eQTLs between tissues we estimated the proportion of replication of eQTLs between all pairs of tissues using the ?1 estimate in the qvalue package [8]. We observed that around 80% of the observed eQTLs active in one tissue also show activity in another tissue (Supplementary Figs 4 and 5). However, the fact that an eQTLs is active in two tissues does not mean that it acts in the same way in both of them. To test if the effect size of eQTLs is the same in two tissues we used a bivariate approach that models pairs of tissues together and compares the beta estimates of the eQTLs in each tissue [9].

When comparing pairs of tissues, we observed that between 32% and 51% of the eQTLs found in both tissues have the same effect size and direction in both tissues while the remaining 49% to 68% eQTLs have tissue specific effects (Figure 1). We also observed that eQTLs active in more than one tissue tend to be closer to the transcription start site (TSS) than tissue specific eQTLs (Supplementary Fig. 6). However, eQTLs further from the TSS have a weaker effect size; therefore these eQTLs are more likely to be misclassified as tissue specific.

**Figure 1.**
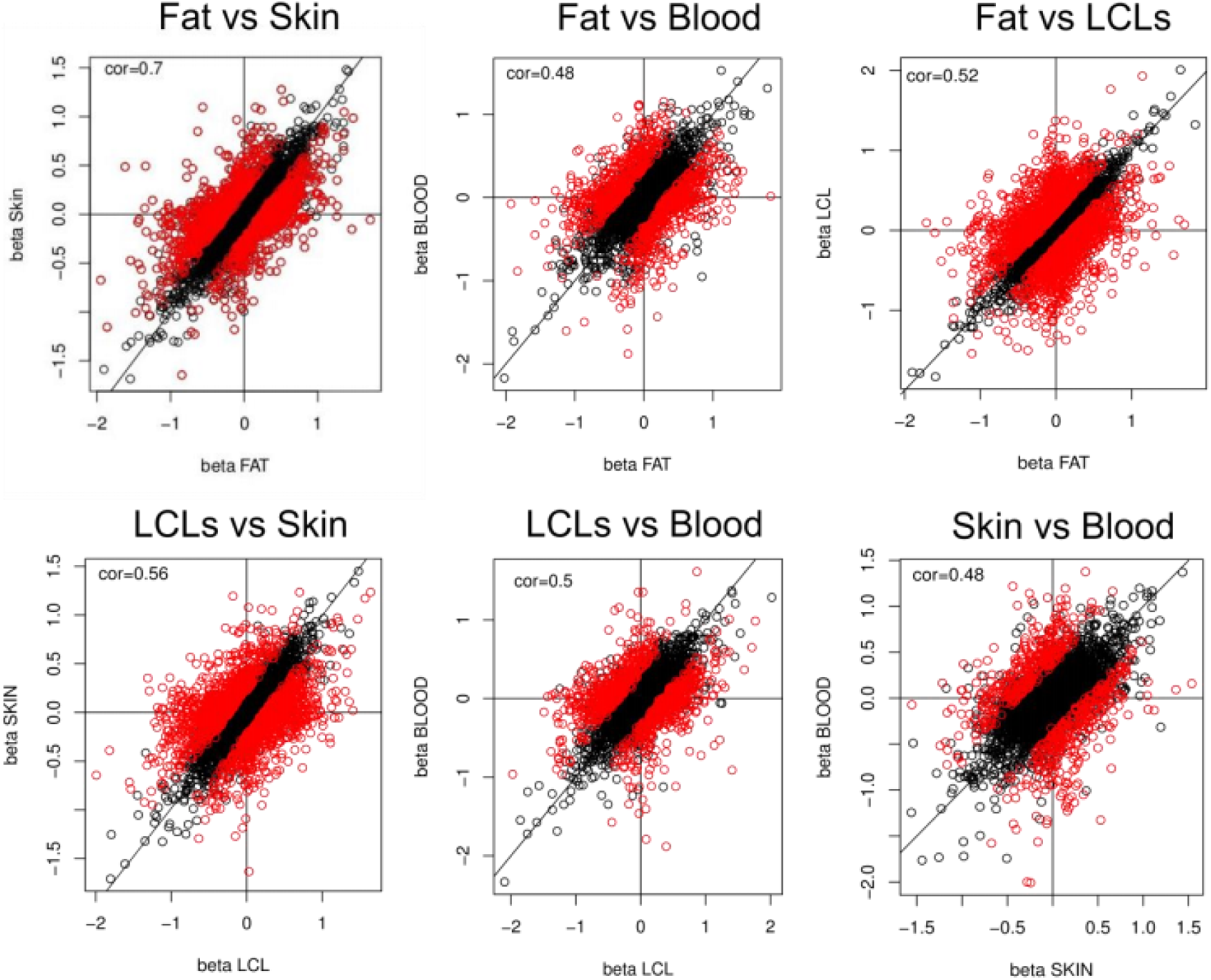
Effect size comparison of cis eQTLs in fat and skin. We compare the direction of effect of eQTLs between fat and skin association. Each plot compares the beta estimated from the SNP-gene associations between two tissues. Red dots identify eQTLs only significant in one tissue.

To understand the biology underlying shared eQTLs relative to distance to the TSS, we looked at the enrichment of eQTLs for each tissue in proximal (2Kb around the TSS) and distal (farther than 2Kb from the TSS) transcription binding sites (TFBS) defined by ENCODE [10]. We found a similar pattern of enrichment in the four tissues for eQTLs falling in proximal TFBS whereas distal eQTLs showed different patterns of enrichment depending on tissue, reinforcing the idea that the biological mechanisms behind the regulation close to the TSS is shared among several tissues while regulatory elements far from the TSS are more tissue specific (Figure 2). Finally, we calculated enrichment of eQTLs in enhancers discovered in several tissues by the FANTOM5 project [11]. We found that the patterns of enrichment in enhancers are tissue specific. eQTLs discovered in blood and LCLs were more enriched in blood cell enhancers whereas eQTLs discovered in skin were more enriched in fibroblast and skin cells enhancers (Figure 2). In conclusion, we observed that genetic regulation in proximal TFBS are shared among tissues while distal TFBS and enhancers act in a more tissue specific manner.

**Figure 2.**
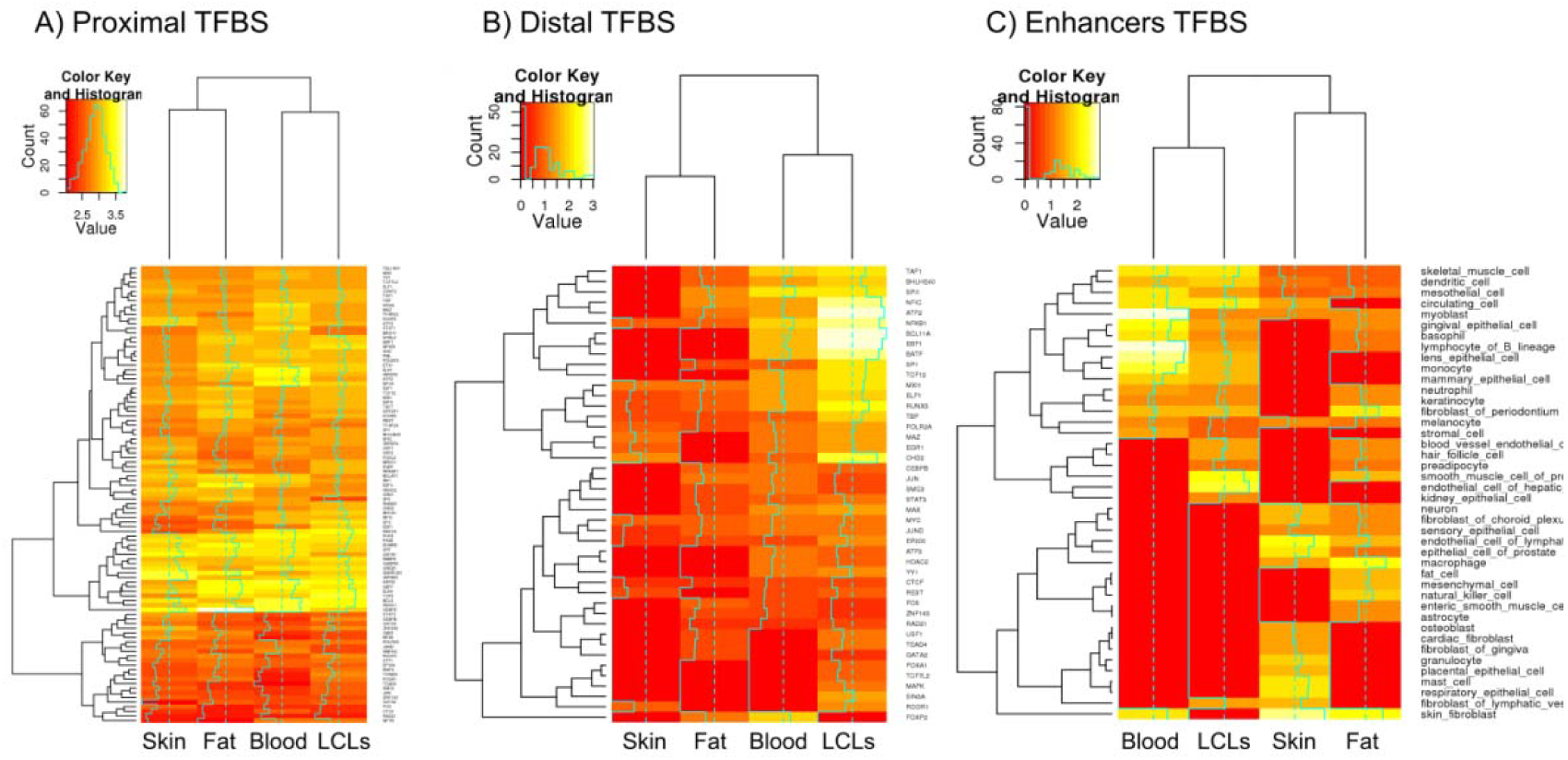
Comparison of the enrichment patterns in fat, skin, LCLS and blood. To understand the biological effects of shared eQTLs, we looked at the enrichment of cis eQTLs for each tissue in a set of markers. A) transcription binding sites (TFBS) defined by ENCODE as proximal TFBS (in 2Kb around the TSS) show similar pattern of enrichment in the four tissues for eQTLs falling in proximal TSS. B) transcription binding sites (TFBS) defined by ENCODE as distal TFBS (farther than 2Kb of the TSS) show a different pattern of enrichment across tissues for eQTLs falling in distal TSS regions. C) enhancers discovered in several tissues by the FANTOM5 project tend to be tissue specific, with cis eQTLs discovered in blood and LCLs more enriched in blood cell enhancers and cis eQTLs discovered in skin more enriched in fibroblast and skin cells enhancers.

The regulatory variants identified by cis-eQTLs analysis represent only a fraction of the influence of genetic variation on gene expression [4]. To quantify the relative contribution of the different genetic and environmental components on gene expression we performed a variance components analysis for each gene separately in each of the 4 tissues using the identity by descent (IBD) information among the twins. We included identified cis-eQTLs, and undetected cis and trans effects as genetic components, as well as shared and individual environmental components in the model. We observed an average heritability of gene expression ranging from 13% to 33%. Of the genetics effects, 53% were trans effects and only 14.5% were due to identified cis eQTLs (Supplementary Figs 7 to 10), with 32.5% due to undetected cis signals. We conclude that most of the genetic variation responsible for differences in expression remains unidentified, meaning that assessing tissue specificity only using identified eQTLs may be misleading. This means that in order to provide an estimate of the shared genetic regulation between tissues it is important to take into account genome-wide genetic effects and not only the known cis-eQTLs.

In order to compare the genetic contribution to gene expression among tissues, we performed a bivariate variance components analysis, considering simultaneously expression of a given gene in a pair of tissues [12]. We produced estimates of genetic and environmental correlations, namely the proportions of the additive genetic and environmental effects that are shared between pairs of tissues.

For genes with heritability larger than 0.2 in both tissues, the mean genetic correlation ranges from 0.04 between fat and blood to 0.17 between fat and skin (Figure 3 and Supplementary Fig. 11); positive correlations indicate shared genetic effects with the same direction. For genes with low heritability, we occasionally observed negative values (Figure 3), that we attribute to poor estimates due to weak effects and small sample size (Supplementary Fig 12). In summary, genes with high heritability in multiple tissues tend to show positive genetic correlation, implying shared genetic effects regulating their expression.

**Figure 3.**
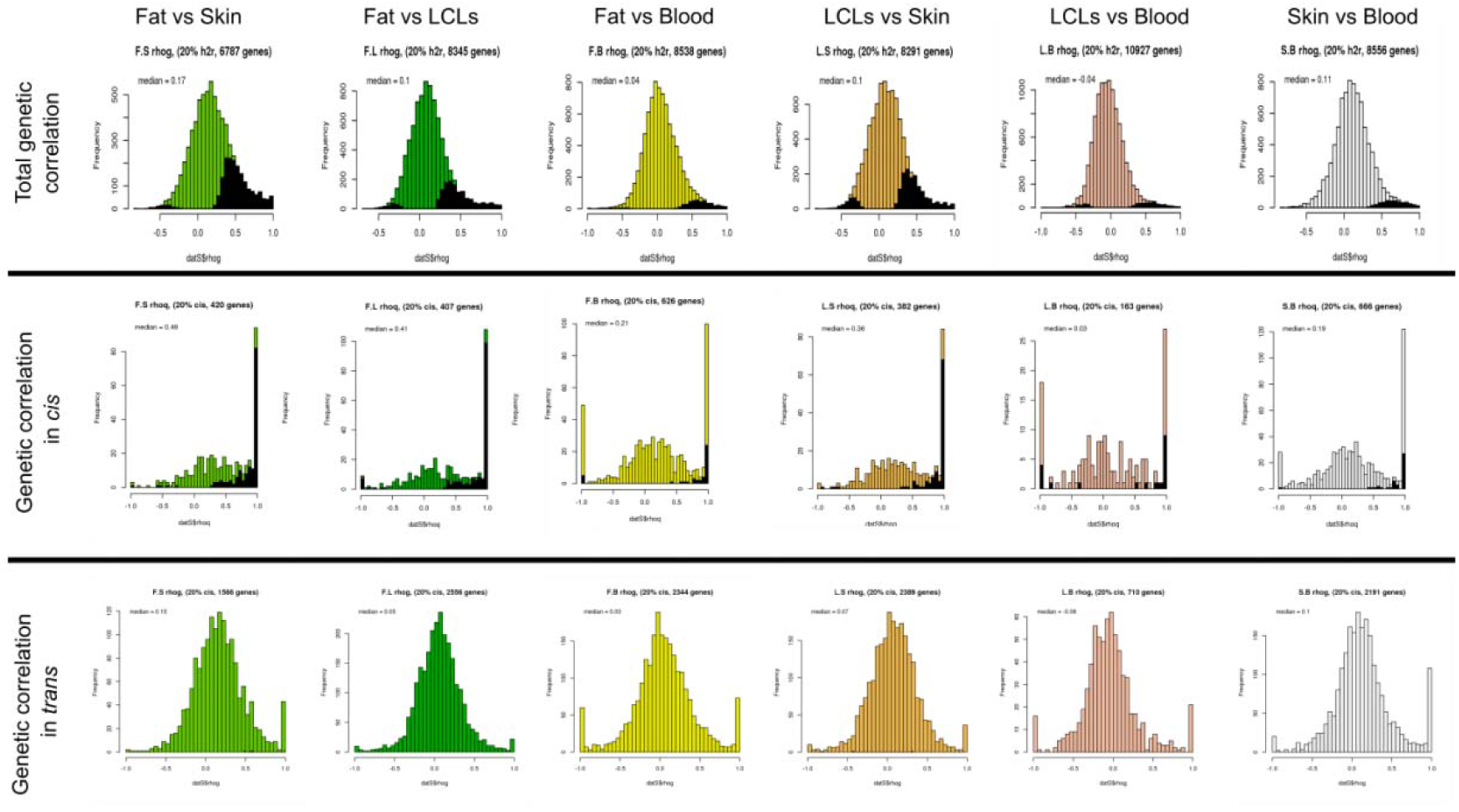
Genetic correlations between tissues. Each column shows results for genetic correlation calculation between two tissues using genes with a total heritability > 0.20 (20%). The first row shows the total gene correlation between tissues, the second row shows the genetic correlation of cis effects and the final row shows the genetic correlation of trans effects.

For most genes, expression has an oligogenic genetic architecture with one or several strong genetic effects in the proximity of the gene (cis effects) and a polygenic effect caused by multiple variants spread along the rest of the genome (trans effects). The genetic correlation measured above is a global measure of the shared genetic effects along the genome, but we are also interested in the differences between cis and trans behavior regarding the shared genetic effects. Previous literature has reported cis effects to have a stronger effect size and are mainly shared across tissues while trans effects tend to be weaker and tissue specific [4] [5] [13] [14]. To obtain separate estimates of the genetic correlation influenced by cis effects and trans effects, we used a bivariate variance components model with three random effects: a cis component, a trans component and a residual component that includes environmental and technical influences (See Methods). We observed clear overrepresentation of genes where the genetic correlation in cis > 0.9, indicating a high degree of sharing of cis effects between tissues (Figure 3). However, for genes with a cis heritability larger than 0.2, the median genetic correlation ranges from 0.03 between LCLS and blood to 0.49 between fat and skin. This indicates that the majority of genetic effects in cis are tissue-specific. As expected, for most of the tissue pairs we found that the distribution of genetic correlations in trans is close to zero (Figure 3). This indicates limited sharing of trans effects, in agreement with previous studies reporting no genetic correlation in trans between fat and blood [13]. However, when considering fat and skin expression, we see a considerable bias towards positive trans correlation, indicating the presence of shared trans effects. In genes with a trans heritability component larger than 20%, the average trans genetic correlation is 0.15, increasing to 0.19 for genes with trans heritability components larger than 30% (Figure 3). This suggests that it would be possible to detect shared trans eQTLs for fat and skin with larger sample sizes.

We then assessed whether the environmental components are shared among tissues. One could hypothesize that all tissues are exposed to the same environment and therefore the non-genetic components of variability are largely shared. We find that the correlation due to non-genetic factors is zero in most of the tissue pairs. Only in fat and skin we observe 454 genes with environmental correlation larger than zero at 5% FDR (Supplementary Figs. 13 to 18). That indicates that non-genetic factors (environment plus technical effects) are mainly tissue specific and highlights very complex processes by which our cells and tissues perceive environmental exposures.

To assess the importance of shared genetic effects for a given tissue, and to place an upper bound on our ability to predict genetic expression in one tissue from another, we can estimate how much of the genetic contribution to a gene in tissue 1 is shared with tissue 2 by multiplying the heritability of the gene in tissue 1 by the genetic correlation between the two tissues. We calculated this magnitude separately for cis and trans heritability. For genes with heritability larger than 0.2, the fraction of the heritability in one tissue that is shared with other tissues ranges from 6% to 35% for the cis component and from 4% to 19% for the trans component (Figure 4). The largest numbers are for fat with skin, sharing 35% of the cis effects and 19% of the trans effects (Figure 4). These numbers are much smaller for other pairs of tissues, especially for the trans component, reflecting once more the fact that trans effects tend to be tissue specific (Figure 4). This result highlights the complexity of regulatory mechanisms between tissues and the high degree of tissue specificity.

**Figure 4.**
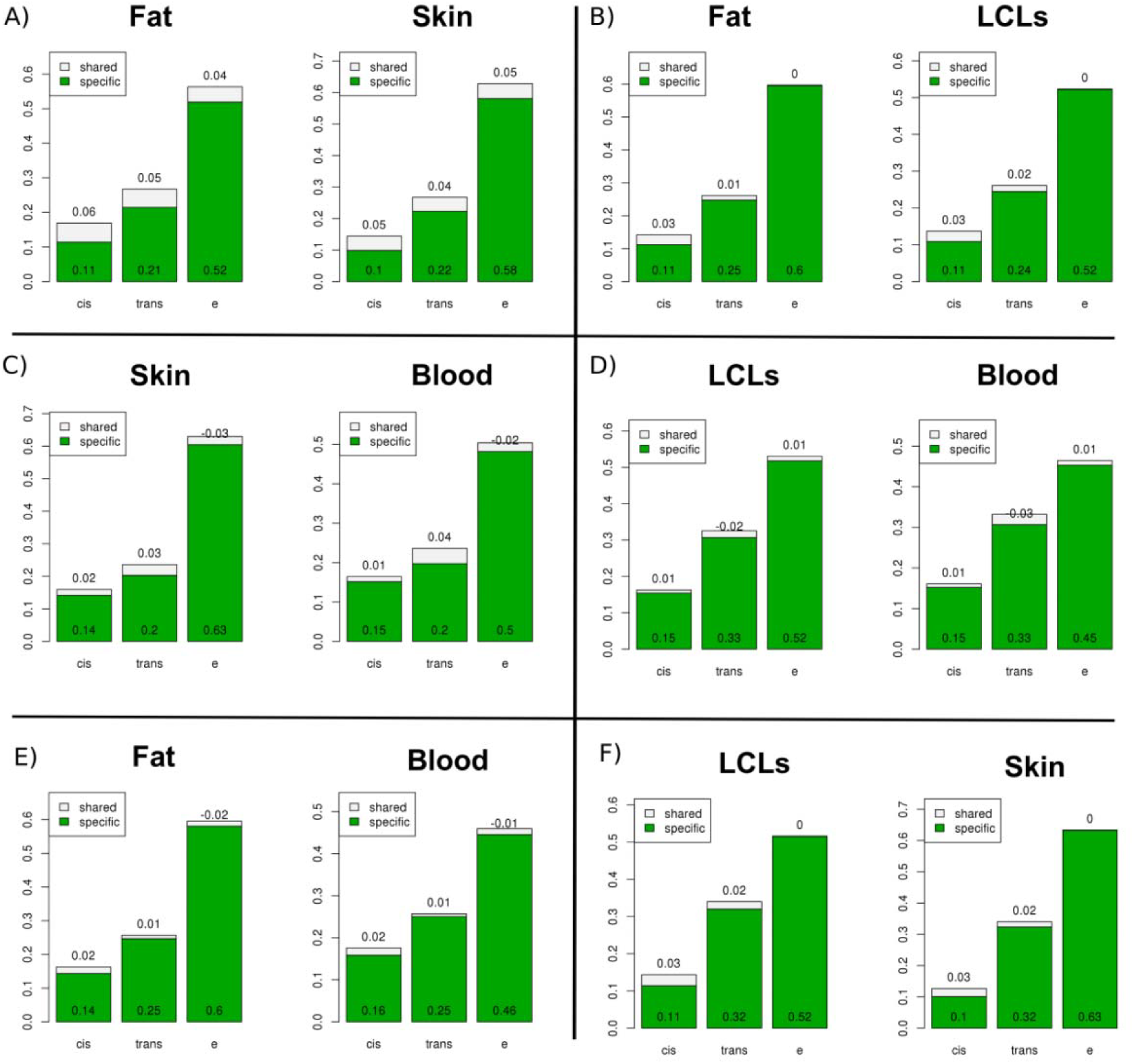
Comparative of the proportion of the genetic effect in between pairs of tissues. We estimate how much of the genetic effects of a gene in tissue 1 is shared with tissue 2 by multiplying the heritability of the gene in tissue 1 by the genetic correlation between the two tissues. We did it separately for cis and trans effects and in genes with heritability values larger than 0.2. The largest numbers are for fat with skin, sharing 35% of the cis effect and 19% of the trans effects (figure A).

In summary, we have presented a systematic genome-wide evaluation of the relative contribution of genetic and non-genetic factors on gene expression across tissues. eQTLs results are frequently used to reveal the biological causes of GWAS signals on complex diseases. Since eQTLs are usually obtained from accessible cell types such as blood or blood cell cultures, it is of special interest to assess how well this information translates to disease relevant tissues [3] [2] [1]. We have demonstrated that common cis eQTLs active in one tissue are likely to be active in other tissues (especially when the eQTLs is close the TSS of the gene of interest), though the effect can be modulated by other factors to produce differing effects. Therefore, the use of functional annotation information in a tissue to interpret the effect of regulatory variants in another tissue is only valid for regulatory variants in the proximity of the TSS of genes with cis-regulatory effects. We have also shown this shared component caused by known eQTLs explains only a minority of the genetic variance, and that genetic variation in the regulation of gene expression is, to a large extent, occurring in trans and is highly tissue specific. In addition, the environmental components of variants are also highly tissue specific suggesting that each tissue and cell type is exposed to different components of the environment and/or it perceives the same environment differently. Overall, our results demonstrate that there is an upper bound to the utility of gene expression in heterologous accessible tissues (e.g. blood) to infer causes of variation in gene expression in inaccessible tissues. This upper bound is less restrictive in biologically interacting tissues such as fat and skin, where there was evidence of shared trans effects. Taken together, these results show the difficulties inherent in predicting and assessing causes of variability of expression of one tissue from another, and highlight the importance and need of studies in many tissues in large samples and environmental contexts to provide a global picture of genetic regulation and its implications on disease at the organismal level.

## Methods

### Sample collection

The study included 856 Caucasian female individuals recruited from the TwinsUK Adult twin registry. Punch biopsies (8mm) were taken from a photo-protected area adjacent and inferior to the umbilicus. Subcutaneous fat tissue was dissected from each biopsy, weighed and immediately stored in liquid nitrogen. Similarly, the remaining skin tissue was weighed and stored in liquid nitrogen. Peripheral blood samples were collected and lymphoblastoid cell lines (LCLs) were generated by Epstein Barr Virus transformation of the B-lymphocyte component by the European Collection of Cell Cultures agency.

The St. Thomas’ Research Ethics Committee (REC) approved on 20th September 2007 the protocol for dissemination of data, including DNA, with the REC reference number RE04/015. On 12th of March of 2008, the St Thomas’ REC confirmed this approval extends to expression data. Volunteers gave informed consent and signed an approved consent form prior to the biopsy procedure. Volunteers were supplied with an appropriate detailed information sheet regarding the research project and biopsy procedure by post prior to attending for the biopsy.

### Genotying and imputation

Samples were genotyped on a combination of the HumanHap300, HumanHap610Q, 1M-Duo and 1.2MDuo 1M Illumnia arrays. Samples were imputed into the 1000 Genomes Phase 1 reference panel (data freeze, 10/11/2010) [15] using IMPUTE2 [16] and filtered (MAF<0.01, IMPUTE info value < 0.8). Only autosomal SNPs were used in the analysis.

### RNA processing

Samples were prepared for sequencing with the Illumina TruSeq sample preparation kit (Illumina, San Diego, CA) according to manufacturer’s instructions and were sequenced on a HiSeq2000 machine. Afterwards, the 49-bp sequenced paired-end reads were mapped to the GRCh37 reference genome [17] with BWA v0.5.9 [18]. We use genes defined as protein coding in the GENCODE 10 annotation [19]. We excluded samples that failed in the library prep or sequence process. We also excluded samples with less than 10 million reads sequenced and mapped to the exons. Finally we excluded samples in which the sequence data did not correspond with the actual genotype data. We ended with 766 samples for fat, 814 for LCLS, 716 for skin and 384 for blood (we had blood samples for only half of the individuals).

## eQTLs discovery

### Exon quantifications

All overlapping exons of a gene were merged into meta-exons with identifier of the form “geneID_start.pos_end.pos”. We counted a read in a meta-exon if either its start or end coordinate overlapped a meta-exon.

### Normalization

All read count quantifications were corrected for variation in sequencing depth between samples by normalizing the reads to the median number of well-mapped reads. We used only exons quantified in more than 90% of the individuals. We removed the effects of technical covariates regressing out the first 50 factors from PEER [20], including BMI and age in the model to preserve important biological sources of variation.

### eQTLs association

Since our data samples are twins, they are not independent observations and we needed to take that into account in our models. We used the two-steps strategy described in Aulchenko et al. [21]. First we kept the residuals of a mixed model that removed the effects of the family structure using the implementation in GenAbel R package. We then transformed those residuals using a rank normal transformation. Finally, we performed a linear regression of the transformed residuals on the SNPs in a 1Mb window around the transcription start site for each gene, using MatrixeQTL R package [22]. We did the association at the exon level and we kept the best association per gene.

### Permutations

We permuted the quantifications of each exon 2000 times, keeping the best p-value per exon from each round. From these data, we adjusted the empirical FDR to 1% according to the most stringent exon of each gene, stratifying the analysis on the number of exons for a given gene.

## Alternative splicing analysis

To calculate the relative quantification of splicing events we used the ALTRANS method that utilizes the paired-end nature of the RNA-seq experiment [7]. It uses the mate pairs, where one mate maps to one exon and the other mate to a different exon, to count “links” between two exons. The first exon in a link is referred to as the “primary exon”. Overlapping exons are grouped into “exon groups” and unique portions of each exon in an exon group are identified, and subsequently used to assign reads to an exon. The raw link counts were normalized with the same method and covariates as described in the “RNA sequencing and quantification” section. The normalized link counts ascertained from unique regions of exons, which can be derived from parts of the linked exons rather than the whole exons, are divided by the probability of observing such a link given the empirically determined insert size distribution for each sample and unique portions of the exons in question, which is referred to as “link coverage”. Finally, the quantitative metric produced is the fraction of one link’s coverage over the sum of the coverages of all the links that the primary exon makes. We calculated this metric in 5’-to-3’ (forward) and 3’-to-5’ (reverse) directions to capture splice acceptor and donor effects respectively. In the association analyses, we only included links where the primary exon’s exon group made at least 10 links in the analyzed direction in at least 80% of the individuals and where the primary exon made at least 5 links in the analyzed direction in at least 30% of the individuals. Furthermore, links with more than 95% non-variable values across were filtered out. The association tests were done as described in the “cis eQTLs association” section.

## IBD calculations

### IBD

We calculated the haplotypes in a 1Mb window around the TSS of each gene and counted the number of haplotype alleles that are shared between the twin pairs at each locus.

## Variance components models (VCM)

All of the variance component analysis were done using gene-based RPKM measures instead of the exon quantifications used in eQTLs analyses. Gene quantifications were attained by summing up all the exon counts of the gene. We only included exons and genes that were expressed in 90% of the samples. Using these counts, reads per kilobase per million reads (RPKM) values were calculated. We removed technical effects by correcting the RPKM measures of each gene for four technical covariates (mean GC content, insert size, primer index used in the library prep and date of library prep) in a linear model.

Variance components models (also called linear mixed models) accommodate the non-independence of family-related individuals and allow the partition of the variance of a quantitative trait (like gene expression of a gene) in several genetic and environmental components. Let’s *Y =* (*y*_1_, *y*_2_) be the phenotype for the individuals in a family (twin pair), we assume normality of *Y* and:

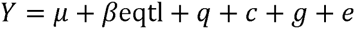

where *μ* is the mean, eqtl is the primary eqtl for the gene, *β* is the regression slope for the eqtl fixed effect, *q* is a random effect capturing the genetic effects in cis (other than the main eqtl), *g* is a random effect capturing the polygenic effects in trans, *c* is a random effect capturing the environment shared between the members of the family and *e* is the residual random effect that includes individual environmental effects. We can express the covariance between relatives as:

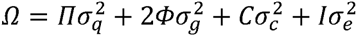

where,

*Π* is the IBD matrix calculated around the TSS of a gene for each pair of twins.

*2Φ* is the matrix of kinship coefficients between pairs of relatives. It is 1 for MZ twin pairs and 1/2 for DZ twin pairs and 0 otherwise.

*C* is the matrix capturing the shared environment between twin pairs. It is 1 for both MZ and DZ twin pairs and 0 otherwise.

*I* is the identity matrix of dimension 2.

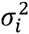 the variance due to genetic effects in cis other than the main eqtl (*q*), genetic effects in trans (*g*), environmental effects shared among the two twins (*c*) and individual environmental effects (*e*).

We estimated the parameters using maximum likelihood methods as implemented in SOLAR [9]. We used the likelihood ration test to test the statistical significance of the parameters.

## Bivariate variance components models

This model is a straightforward extension of the univariate model described above. Let *X* = (*x*_1_,*x*_2_)′ and *Y* = (*y*_1_,*y*_2_)′ be the twin pair trait vectors for two phenotypes. We assume that *X* and *Y* are normally distributed as in the univariate case:

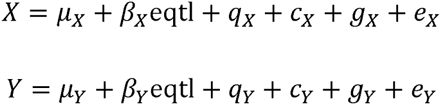

and have covariance matrices:

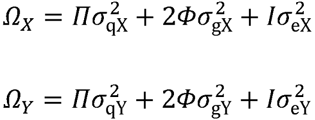

Note that in the bivariate case we did not include the shared environment component.

Then, we can express the bivariate phenotype as

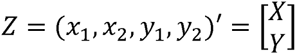

The covariance matrix for Z has the partition structure,

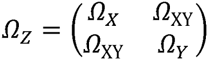

Where Ω_*X*_ and Ω_*Y*_ are the univariate covariance matrices described above and the matrix Ω_*XY*_ = Ω_*YX*_ of cross covariances is given by,

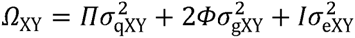

We can reparametrize the covariances in terms of correlations by writing,

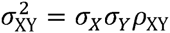

Where *ρ*_XY_ is the correlation between traits *X* and *Y.* The complete covariance matrix for Z can be written,

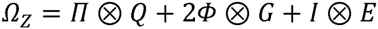

Where ⊗ is the Kronecker product operator and *n* is the size of the pedigree. For 2 traits is a twin pair (2 individuals), matrices Q, G and E are 2 × 2 matrices, *Π, Φ* and *I* are 2 × 2 and Ω_*Z*_ is 4 × 4. The matrices Q, G, and E of QTL-specific, polygenic, and environmental variance components respectively, each have the partition form,

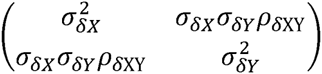

Where *ρ*_δXY_ is the correlation between *X* and *Y* due to the effect of *δ*, and *δ* is q, g or e.

We have then a model with 11 parameters: the averages of the two traits (μ_*X*_and μ_*Y*_), the 3 variance components of the two traits 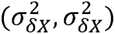 for *δ* in *q, g* or *e* and the 3 correlations for cis, trans and environment components (*ρ*_qXY_,*ρ*_gXY_, *ρ*_eXY_).

In the same way that in the univariate case, to estimate the parameters we used maximum likelihood methods as implemented in SOLAR [9]. We used likelihood ratio tests to establish the statistical significance of the parameters.

## Bivariate model simulation

We tested the performance of the bivariate VCM in our sample by simulating bivariate phenotypes under several models, always with our actual pedigrees,

Case 1: two traits with heritability of 0.2 and genetic correlation equal to 1

Case 2: two traits with heritability of 0.2 and genetic correlation equal to 0.5

Case 3: two traits with heritability of 0.2 and genetic correlation equal to 0

Case 4: two traits with heritability of 0 so that genetic correlation is not well defined.

We performed 100 simulations of each model using the command simqtl in SOLAR and maximized the bivariate model for all the simulations. The results show that we get unbiased estimated of the genetic correlations in all the cases (Supplementary Figure 12). For the first case (rhog=1) we got an estimate of 1 for most of the simulated replicates, with an average of 0.9. For the second case (rhog=0.5) we got an average of 0.5 but with considerable variation. Than implies that, with this sample size, it is going to be difficult to get statically significant estimates of rhog, but still, we do not have biases results. For the third case (rhog=0) we got a distribution of genetic correlations estimates symmetrical and centered on zero. For the fourth case (not heritable traits) we got a distribution with the average close to zero but two unwanted peaks in the values 1 and −1. Since the genetic correlation is not defined when the traits are not heritable, the likelihood is not affected by the value of rhog and, for some reason, the maximization procedure tends to end with values of rhog in the extreme of the parameter space (1 and −1). Due to the lack of precision of the estimates, we observe in all cases many genes with negative genetic correlation. In theory, a negative genetic correlation means that the two traits have common genetic effects that affect both traits in different directions. However in the context of our analysis these negative values are mainly a consequence of the variance in the estimates due the limited sample size. Although we cannot rule out the possibility that for some genes a negative estimate of the genetic correlation could reflect a true biological fact, it is impossible for us to tell these case from the more common negative genetic correlations due to lack of precision in the estimates. In summary, average estimates of rhog in our data should be close to truth even that the individual estimates per gene can be far from the real values. For some analysis we give results only for genes that have heritability larger than 0.2 in both tissues so that we avoid cases where rhog is not defined.

## Enrichment Analysis

Given a genomic functional category (like enhancer or TFBS) we calculate the enrichment of eQTLs in this category by comparing the number of eQTLs that fall into this category to the number of comparable random SNPs that fall in the same category. We generated the comparable set of random SNPs choosing random SNPs that match in allele frequency and distance to the TSS of the gene with the real eQTLs. We generated a maximum of 10 matched random SNPs for each eQTLs and calculated odds ratio and a p-value using Fisher exact test.

In our analysis we used two sets of functional annotation information. We used FAMTOM5 annotation of enhancers in a tissue and ENCODE annotation of TFBS. We divided the TFBS in two sets, proximal TFBS are those located less than 2Kb from the TSS of the gene and distal TBFS are the rest.

